# Sentinel-2 based estimates of rangeland fractional cover and canopy gap class for the western United States

**DOI:** 10.1101/2025.03.13.643073

**Authors:** Brady W Allred, Sarah E. McCord, Timothy J. Assal, Brandon T. Bestelmeyer, Chad S. Boyd, Alexander C. Brooks, Samantha M. Cady, Michael C. Duniway, Samuel D. Fuhlendorf, Shane A. Green, Georgia R. Harrison, Eric R. Jensen, Emily J. Kachergis, Anna C. Knight, Chloe M. Mattilio, Brian A. Mealor, David E. Naugle, Dylan O’Leary, Peter J. Olsoy, Erika S. Peirce, Jason R. Reinhardt, Robert K. Shriver, Joseph T. Smith, Jason D. Tack, Ashley M. Tanner, Evan P. Tanner, Dirac Twidwell, Nicholas P. Webb, Scott L. Morford

**Affiliations:** Numerical Terradynamic Simulation Group, University of Montana, Missoula, MT, USA; Jornada Experimental Range, USDA Agricultural Research Service, Las Cruces, NM, USA; Bureau of Land Management, National Operations Center, Denver, CO, USA; Eastern Oregon Agricultural Research Center, USDA Agricultural Research Service, Burns, OR, USA; Desert Research Institute, Reno, NV, USA; Department of Agronomy and Horticulture, University of Nebraska–Lincoln, Lincoln, NE, USA; U.S. Geological Survey, Southwest Biological Science Center, Moab, UT, USA; Natural Resource Ecology and Management, Oklahoma State University, Stillwater, OK, USA; USDA Natural Resources Conservation Service, Central National Technology Support Center, Ft. Worth, TX, USA; University of Wyoming Sheridan Research and Extension Center, Institute for Managing Annual Grasses Invading Natural Ecosystems, Sheridan, WY, USA; W.A. Franke College of Forestry and Conservation, University of Montana, Missoula, MT, USA; Institute for Natural Resources, Oregon State University, Corvallis, OR, USA; Rangeland Resources and Systems Research Unit, USDA Agricultural Research Service, Fort Collins, CO, USA; USDA Forest Service, Rocky Mountain Research Station, Moscow, ID, USA; Department of Natural Resources and Environmental Science, University of Nevada, Reno, NV, USA; US Fish and Wildlife Service, Habitat and Population Evaluation Team, Missoula, MT, USA; Caesar Kleberg Wildlife Research Institute, Texas A&M University-Kingsville, Kingsville, TX, USA

## Abstract

Rangelands are extensive ecosystems, providing important ecosystem services while undergoing continuous change. As a result, improved monitoring technologies can help better characterize vegetation change. Satellite remote sensing has proven effective in this regard, tracking vegetation dynamics at broad and fine scales. We leveraged the spatial, spectral, and temporal resolution of Sentinel-2 satellites to estimate fractional cover and canopy gap across rangelands of the western United States. We produced annual, 10 m spatial resolution estimates of fractional cover and canopy gap size class for years 2018 to 2024. Fractional cover estimates include that of common plant functional types (annual forb and grass, bareground, littler, perennial forb and grass, shrub, tree) and select genera (including invasive annual grass species, pinyon-juniper species, and sagebrush species); canopy gap size classes include gap sizes 25 to 50, 51 to 100, 101 to 200, and greater than 200 cm. We make these data available as Cloud Optimized GeoTIFFs, organized as 75×75 km tiles covering the 17 western states of the United States.

## Background and Summary

Rangelands are diverse, heterogeneous ecosystems that contain a variety of plant communities, including grasslands, shrublands, woodlands, and savannas. Covering approximately half the earth’s land surface^1^, rangelands support a wide array of biodiversity and provide important ecosystem services. Due to their vast coverage and use by humans, monitoring rangeland conditions has been a primary objective for decades^2^. Since the beginning of satellite remote sensing, scientists and engineers have actively pursued mapping and quantifying rangelands at broad scales^3–6^. In recent years, research has focused on estimating fractional cover, i.e., estimating the proportion of an area (commonly the area of a pixel) covered by vegetation or other land cover types^7,8^. Compared to traditional categorical or thematic vegetation classifications, fractional cover estimates better represent the sub-pixel complexity and landscape heterogeneity that dominate rangelands and have become a standard indicator for monitoring ecological state change and the effectiveness of management strategies^9^.

Decomposition and unmixing methods have historically and successfully produced fractional cover estimates of general cover classes, including photosynthetically active and inactive vegetation^10–13^. The increased collection of vegetation data across broad areas–particularly in the United States–resulted in the ability to empirically model fractional cover and other indicators, such as plant invasion and aeolian erosion^14,15^. Fractional cover estimates may be aggregated to specific plant functional types/groups, genera, or remain at the individual species level^16–18^. While the use of satellite imagery is widespread and permits estimates across spatiotemporal scales, aerial imagery can also be used to produce finer resolution cover estimates^19,20^, albeit with limited spatial, temporal, and spectral scope. Empirical methods vary in their algorithmic approach (linear models, support vector machines, etc.) and use of satellite data (single images, full temporal sequences, etc.). Among algorithms, regression trees and neural networks are frequently employed; tree based methods are valued for their ease of use and ability to handle collinearity and non-linear interactions among variables, while neural networks excel at learning complex patterns^21,22^.

At broad extents, rangeland fractional cover estimates have historically been derived from the MODIS and Landsat satellite collections^13,17^. While the images from these collections are spatially coarser than newer sensors (250 m and 30 m spatial resolution, respectively), their advantage is an unmatched historical record that permits unique time series analysis^15,23,24^.

Newer satellites and sensors, however, offer improvements that enable the detection of more subtle changes in rangeland condition and facilitate the exploration of new research and management questions. The Sentinel-2 mission currently consists of three satellites (2A launched June 2015, 2B launched March 2017; 2C launched September 2024 and is planned to replace 2A), producing optical imagery with an increased number of spectral bands, at a finer spatial resolution of 10 to 60 m, and at a shorter nominal revisit time of 10 days. These improved sensor characteristics provide distinct advantages for rangeland mapping. Finer spatial resolution reduces signal mixing from heterogeneous vegetation, enabling the detection of smaller features. Temporally, quicker revisit times (relative to Landsat satellites) may distinguish unique phenological signatures of different plant functional types. These improvements provide more precise spatial, temporal, and spectral information, ultimately leading to more accurate fractional cover estimates.

While fractional cover provides valuable information regarding rangeland vegetation, it does not fully capture the spatial arrangement of plants. Canopy gap, which quantifies the size and distribution of openings between plant canopies, is an important indicator for many ecosystem processes, including wind erosion, water infiltration, and wildlife habitat^25,26^. Estimating canopy gap distributions alongside fractional cover of plant functional types may improve the characterization of rangeland vegetation structure and increase the value of these data for understanding and responding to structural ecosystem change.

We explored the use of Sentinel-2 satellites in estimating fractional cover and canopy gap across the United States. We constructed a temporal one-dimensional convolutional neural network to concurrently estimate the fractional cover of common plant functional types, select genera, and canopy gap size class. Using this model, we produced annual estimates characterizing rangeland vegetation cover and canopy gap across the17 western states of the United States at 10 m spatial resolution for the years 2018 through 2024.

### Methods

#### Vegetation data

We used field data collected by various programs across the conterminous United States (CONUS; Table 1) to train a model capable of estimating fractional cover and canopy gap size classes. Vegetation was sampled on private and public rangelands using line point intercept (LPI) and canopy gap methods in 47 states within CONUS. As field data were collected by separate monitoring programs, methodology differed slightly among collections (e.g., transect length and placement, time of sampling, etc.); methods generally followed protocols established by Herrick et al.^27^ and quality assurance procedures laid out in McCord et al.^28,29^. Data collectors were trained with a rigorous U.S. federal agency training curriculum^30–32^. Field data were aggregated and harmonized as part of the Landscape Data Commons using the R package *terradactyl*^29,33^. Species cover, litter cover, and bare soil were calculated using LPI first hit and aggregated into various plant functional types or genera (Table 2; Figure 1). Canopy gaps were measured and aggregated into classes (Table 2; Figure 2). We limited vegetation data to years 2018 through 2024 (Figure 3), corresponding with the bulk of the Sentinel-2 satellite record in the United States. We only used field samples in which vegetation cover and canopy gap measurements were available, resulting in 47,833 field data samples for model development.

**Figure 1.**
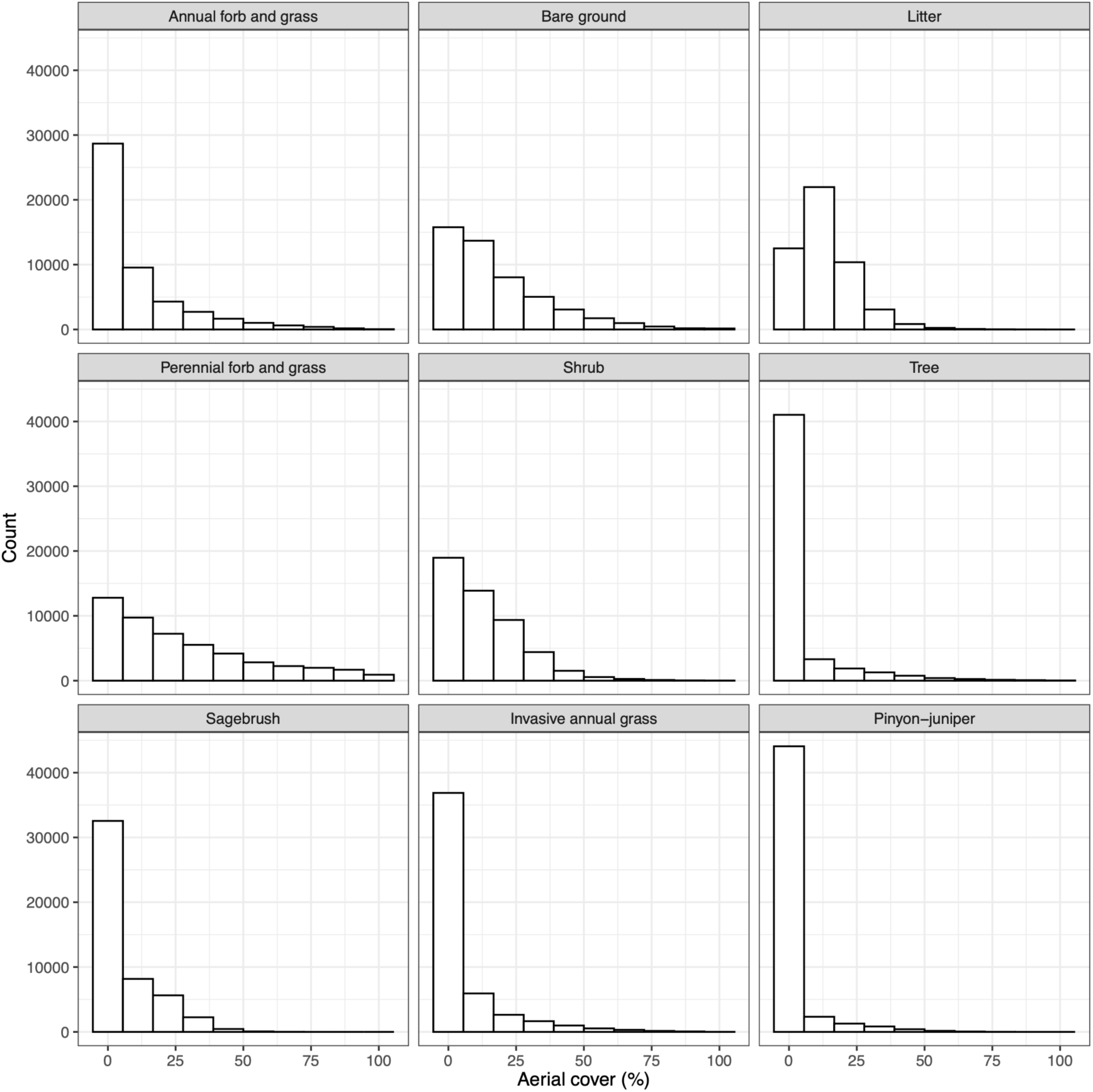
Frequency of cover measurements for the vegetation field data. Vegetation categories are described in Table 2.

**Figure 2.**
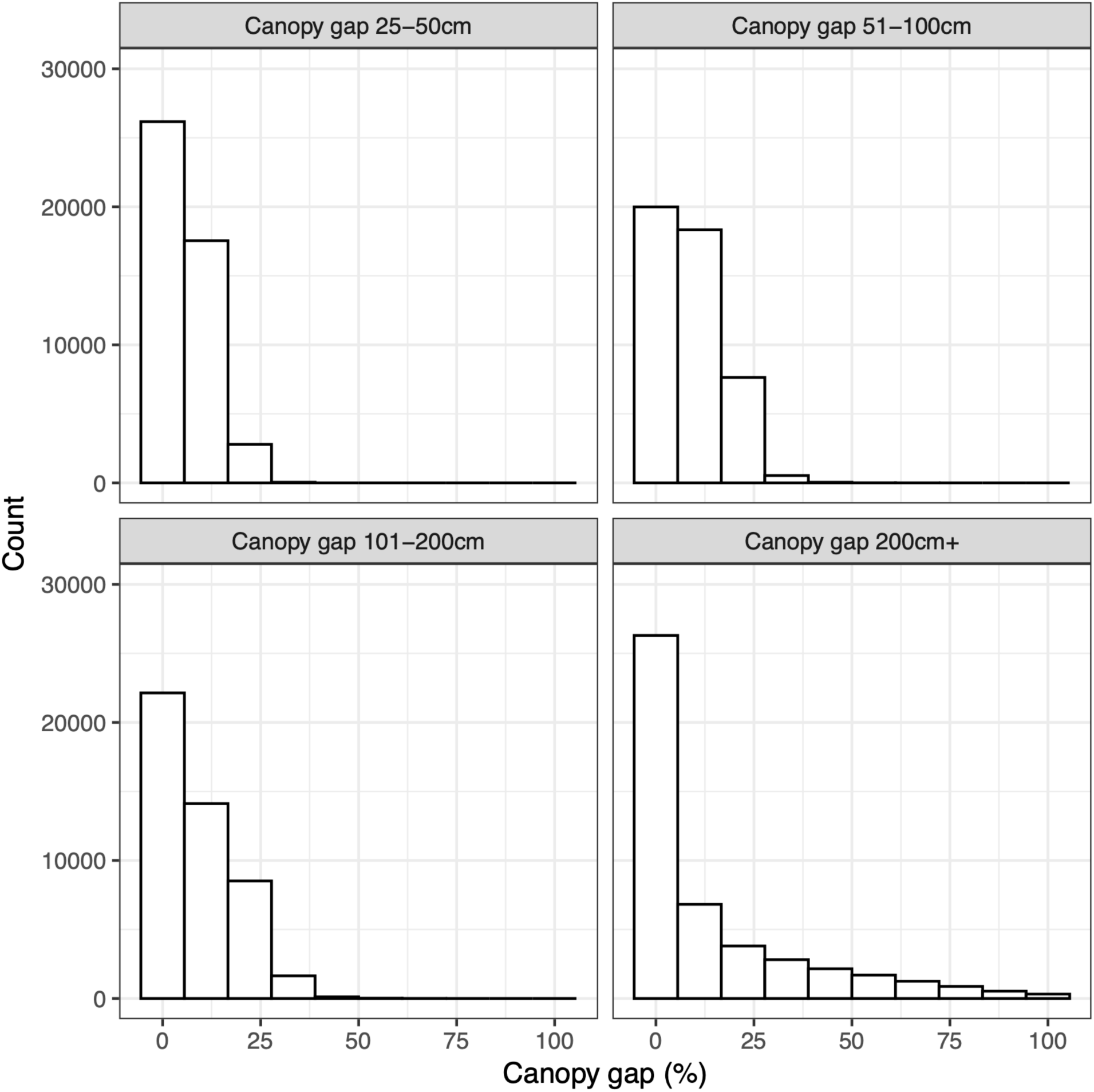
Frequency of canopy gap measurements for the vegetation field data. Canopy gap classes are described in Table 2.

**Figure 3.**
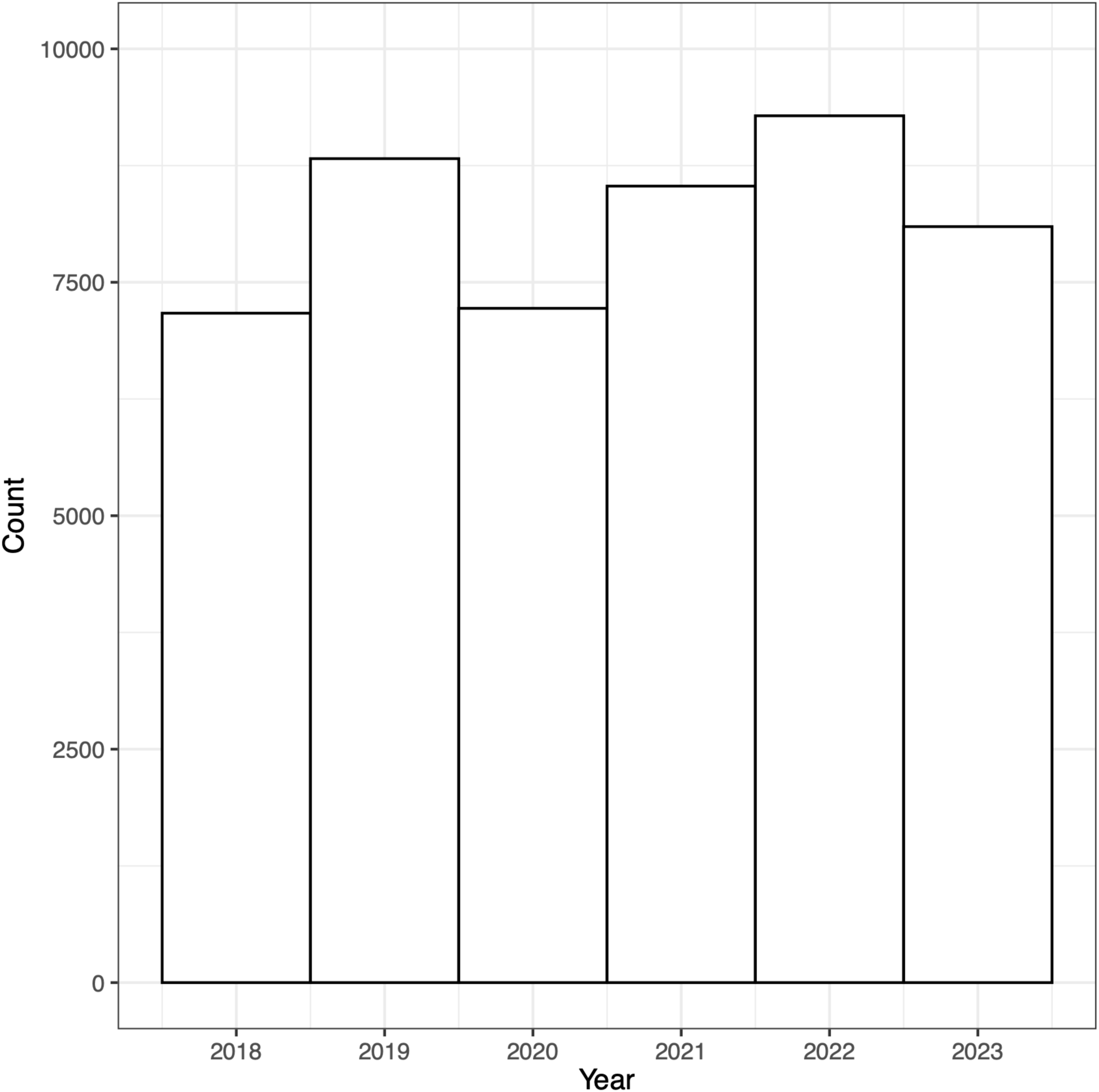
Temporal distribution of sampling plots from the vegetation field data used in model training.

**Table 1.** Data used for model training. Vegetation data were collected using line point intercept methods and span private and public rangelands across the conterminous United States. Data were obtained from the Landscape Data Commons^33^. Additional citations for each data source are provided in the table, where available.

| **Program / Source** | **Number of samples** | **Years** |
| --- | --- | --- |
| Bureau of Land Management Assessment, Inventory, and Monitoring<sup>60</sup> | 39,024 | 2018 - 2023 |
| Bureau of Land Management Assessment, Inventory, and Monitoring - Riparian and Wetland<sup>61</sup> | 407 | 2022 |
| Bureau of Land Management Beartrap RX Burn Monitoring Data | 6 | 2020 - 2023 |
| United States Forest Service Crooked River National Grassland Ecological Site Descriptions | 19 | 2018 - 2022 |
| Jornada Experimental Range | 200 | 2028 - 2022 |
| Nevada Department of Wildlife | 304 | 2018 - 2021 |
| National Park Service Inventory and Monitoring<sup>62</sup> | 224 | 2018 - 2020 |
| Natural Resources Conservation Service National Resources Inventory<sup>63</sup> | 7,318 | 2018 - 2023 |
| National Wind Erosion Research Network<sup>64</sup> | 109 | 2018 - 2022 |
| University of Idaho Rinker Rock Creek Ranch | 58 | 2021 - 2022 |
| United States Geological Survey Post Fire Paired Plots<sup>65</sup> | 164 | 2021 - 2022 |
| National Agricultural Statistics Survey Cropland Data Layer<sup>35</sup> | 9,938 | 2018 - 2022 |

**Table 2.**
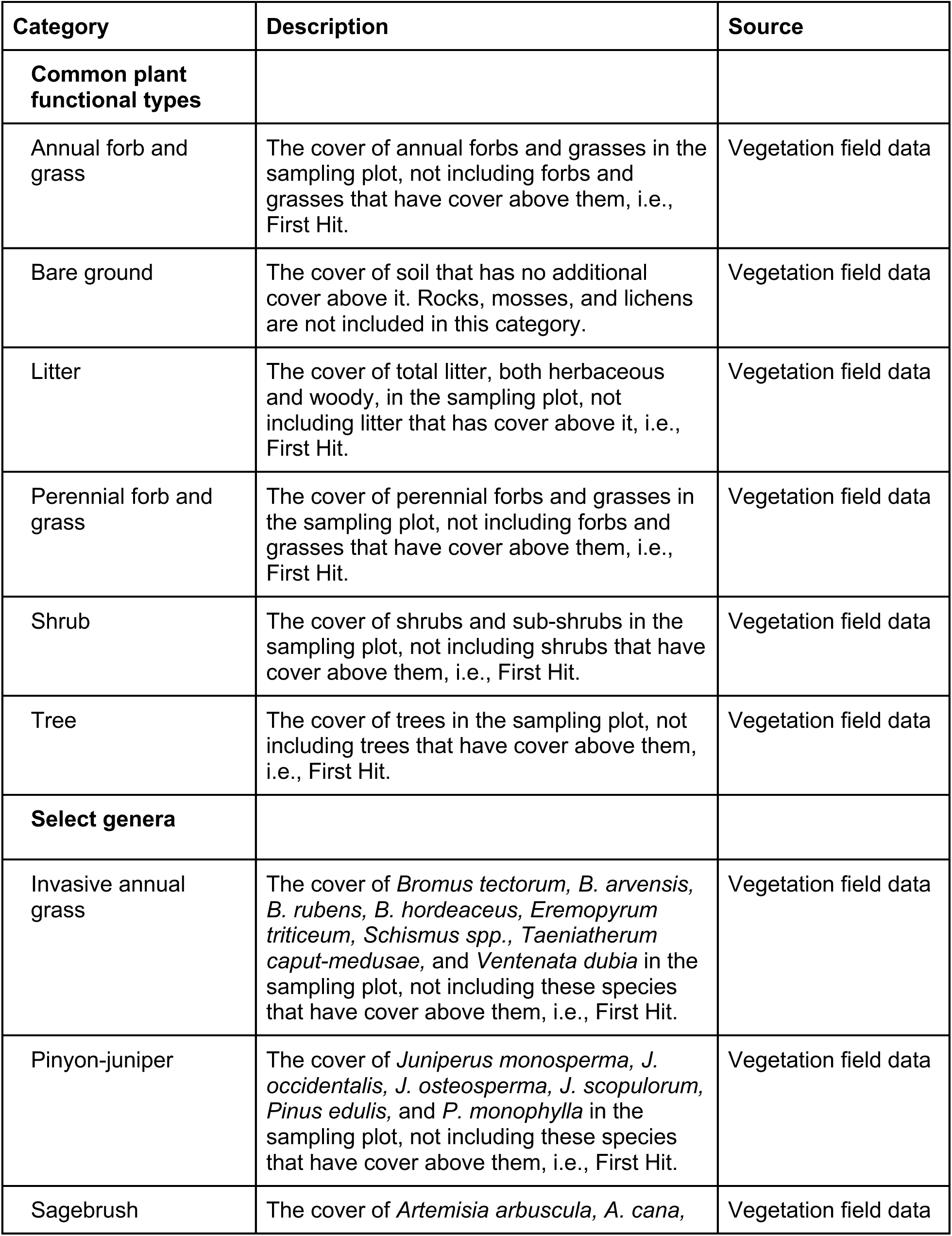
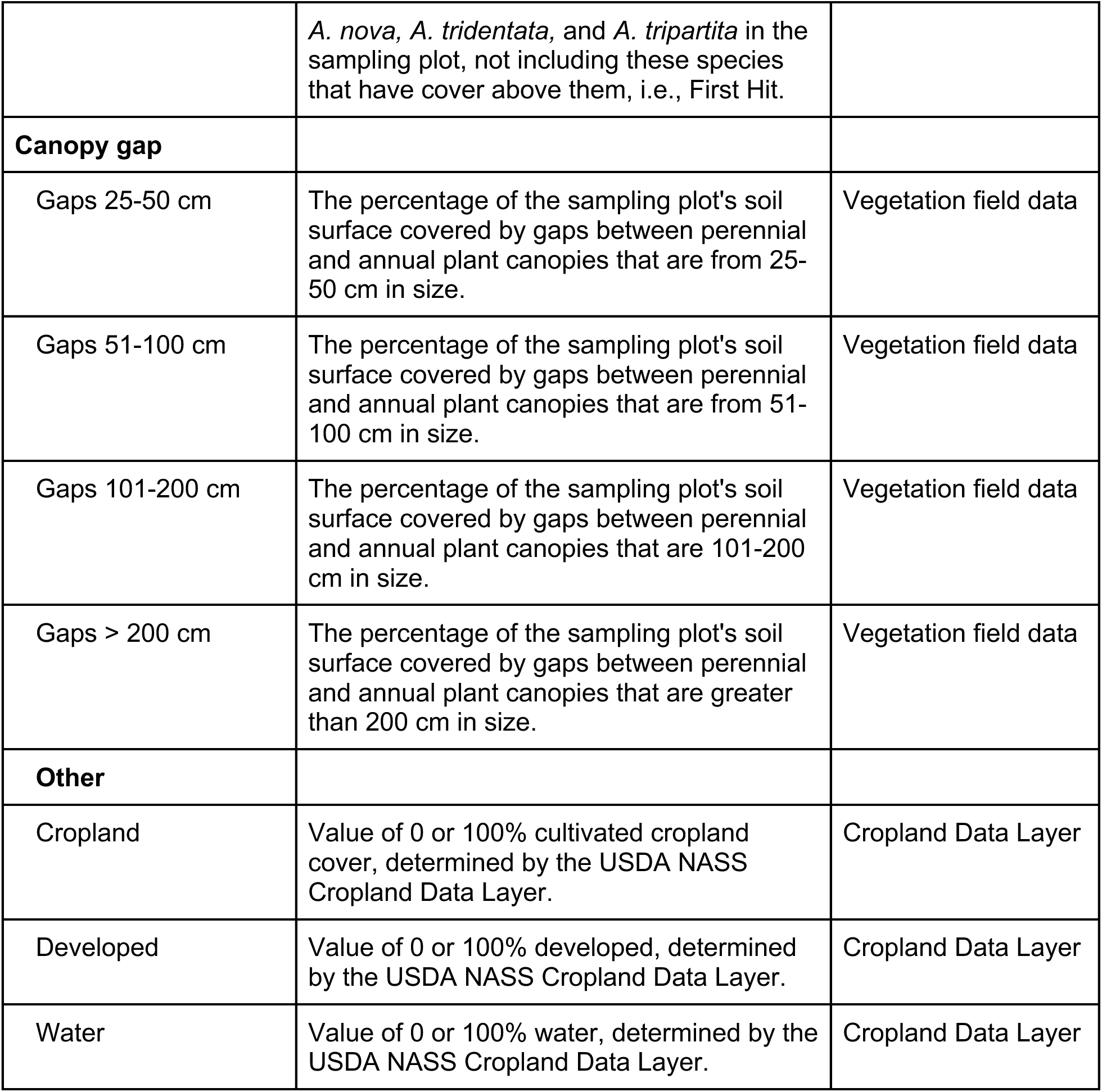
Categories and descriptions of fractional cover predictions and data sources.

| Category | Description | Source |
| --- | --- | --- |
| **Common plant functional types** |  |  |
| Annual forb and grass | The cover of annual forbs and grasses in the sampling plot, not including forbs and grasses that have cover above them, i.e., First Hit. | Vegetation field data |
| Bare ground | The cover of soil that has no additional cover above it. Rocks, mosses, and lichens are not included in this category. | Vegetation field data |
| Litter | The cover of total litter, both herbaceous and woody, in the sampling plot, not including litter that has cover above it, i.e., First Hit. | Vegetation field data |
| Perennial forb and grass | The cover of perennial forbs and grasses in the sampling plot, not including forbs and grasses that have cover above them, i.e., First Hit. | Vegetation field data |
| Shrub | The cover of shrubs and sub-shrubs in the sampling plot, not including shrubs that have cover above them, i.e., First Hit. | Vegetation field data |
| Tree | The cover of trees in the sampling plot, not including trees that have cover above them, i.e., First Hit. | Vegetation field data |
| **Select genera** |  |  |
| Invasive annual grass | The cover of *Bromus tectorum*, *B. arvensis*, *B. rubens*, *B. hordeaceus*, *Eremopyrum triticeum*, *Schismus spp.*, *Taeniatherum caput-medusae*, and *Ventenata dubia* in the sampling plot, not including these species that have cover above them, i.e., First Hit. | Vegetation field data |
| Pinyon-juniper | The cover of *Juniperus monosperma*, *J. occidentalis*, *J. osteosperma*, *J. scopulorum*, *Pinus edulis*, and *P. monophylla* in the sampling plot, not including these species that have cover above them, i.e., First Hit. | Vegetation field data |
| Sagebrush | The cover of *Artemisia arbuscula*, *A. cana*, | Vegetation field data |
|  | *A. nova*, *A. tridentata*, and *A. tripartita* in the sampling plot, not including these species that have cover above them, i.e., First Hit. |  |
| **Canopy gap** |  |  |
| Gaps 25-50 cm | The percentage of the sampling plot's soil surface covered by gaps between perennial and annual plant canopies that are from 25-50 cm in size. | Vegetation field data |
| Gaps 51-100 cm | The percentage of the sampling plot's soil surface covered by gaps between perennial and annual plant canopies that are from 51-100 cm in size. | Vegetation field data |
| Gaps 101-200 cm | The percentage of the sampling plot's soil surface covered by gaps between perennial and annual plant canopies that are 101-200 cm in size. | Vegetation field data |
| Gaps > 200 cm | The percentage of the sampling plot's soil surface covered by gaps between perennial and annual plant canopies that are greater than 200 cm in size. | Vegetation field data |
| **Other** |  |  |
| Cropland | Value of 0 or 100% cultivated cropland cover, determined by the USDA NASS Cropland Data Layer. | Cropland Data Layer |
| Developed | Value of 0 or 100% developed, determined by the USDA NASS Cropland Data Layer. | Cropland Data Layer |
| Water | Value of 0 or 100% water, determined by the USDA NASS Cropland Data Layer. | Cropland Data Layer |

We divided data into training, validation, and testing sets (85%, 5%, and 10%, respectively). The training set was used to fit the model, the validation set to tune hyperparameters and prevent over-fitting during training, and the testing set for the final evaluation of model performance. To ensure independence of our testing data and to mitigate spatial autocorrelation, we used the Generalized Random Tessellation Stratified algorithm in the spsurvey R package^34^ for the selection of our testing set. GRTS is a probabilistic sampling method that maximizes the separation between testing points, producing a spatially balanced sample and avoiding the clustering that may occur with simple random sampling. For each temporal stratum (year of data collection), this process resulted in a test set where points were well distributed among the remaining training and validation points across the United States. The testing set was selected first; afterwards, the training and validation sets were selected.

We aimed for our model to provide estimates in naturally vegetated areas only, excluding areas that were cropland, developed, or water. While this can be accomplished after the fact through pixel masking, we wanted our model to be independent and self-reliant. Ideally, our rangeland dataset would be combined with a cover dataset representing these additional types (i.e., croplands, developed, and water) for our time period and spatial resolution; however, no such dataset exists. To substitute, we used additional data from the Cropland Data Layer^35^, a medium-resolution (30 m) classification dataset that covers the United States. Stratifying across years 2018 through 2022, we randomly selected point locations within CONUS that were classified as cultivated, developed, or open water. To match the continuous measurements of the vegetation data, each location was given a value of 100 according to its classification (i.e., cultivated croplands, development, or water) and a value of 0 for all other groups. While simplistic, the inclusion of these data was to inform the model what is not rangeland, not to accurately estimate fractional cover in cropland, developed, or water areas. This additional dataset was separated into training, validation, and testing (85%, 5%, and 10%, respectively); we used a total of 9,938 additional samples and combined them with the vegetation data.

#### Satellite data

We used Sentinel-2 (satellites 2A and 2B) top of atmosphere reflectance as satellite input. While surface reflectance is commonly used to minimize atmospheric effects, top of atmosphere reflectance provided a greater number of images (available within Google Earth Engine) that intersected CONUS, allowing for a longer time period of satellite data to be used for both model training and inference. Although atmospheric effects can introduce inter-scene variability, our model is trained on a diverse amount of spatiotemporal data, and is sufficiently robust to learn the consistent, underlying signals from top of atmosphere reflectance (refer to *Model performance*). We follow previous modeling efforts that have successfully used top of atmosphere reflectance^36,37^ for dataset development, most notably Dynamic World^38^.

We masked pixels with a Cloud Score+^39^ usability value less than 0.65. For a given calendar year, we created 29 sequential 10-day timesteps, beginning on day of year 041 and ending on day 330, to best capture the growing season and to limit data processing. For each timestep, we calculated the median reflectance value for the visible (bands 2-4), near infrared (bands 5-8; the near infrared bands provide greater spectral capacity compared to Landsat sensors), and short-wave infrared (bands 11-12) bands. Missing reflectance values within a timestep (due to clouds or missing data) were filled using a forward-fill approach (i.e., carrying the value from the previous timestep forward). We chose this method over interpolation and gap filling as it is computationally efficient and scalable for a dataset of this magnitude. Across all samples and time steps, 12.9% of values were missing and were forward filled, with most filling occurring in the earlier and later times of the year (Figure 4). Reflectance values were natural log normalized to reduce skewness in distribution. We also calculated the normalized difference vegetation index (NDVI), a measure of vegetation greenness, calculated as (B8 - B4) / (B8 + B4)^40^; and the normalized burn ratio two (NBR2), an index sensitive to soil and plant water content, calculated as (B8 - B12) / (B8 + B12)^41^. Sentinel-2 bands at resolutions greater than 10 m were resampled to 10 m using nearest neighbor. Spatial location (X,Y coordinates) was calculated in an equal area coordinate reference system (EPSG:5070) and normalized between zero and one using the maximum and minimum coordinates of CONUS. All image processing was performed in Google Earth Engine^42^.

**Figure 4.**
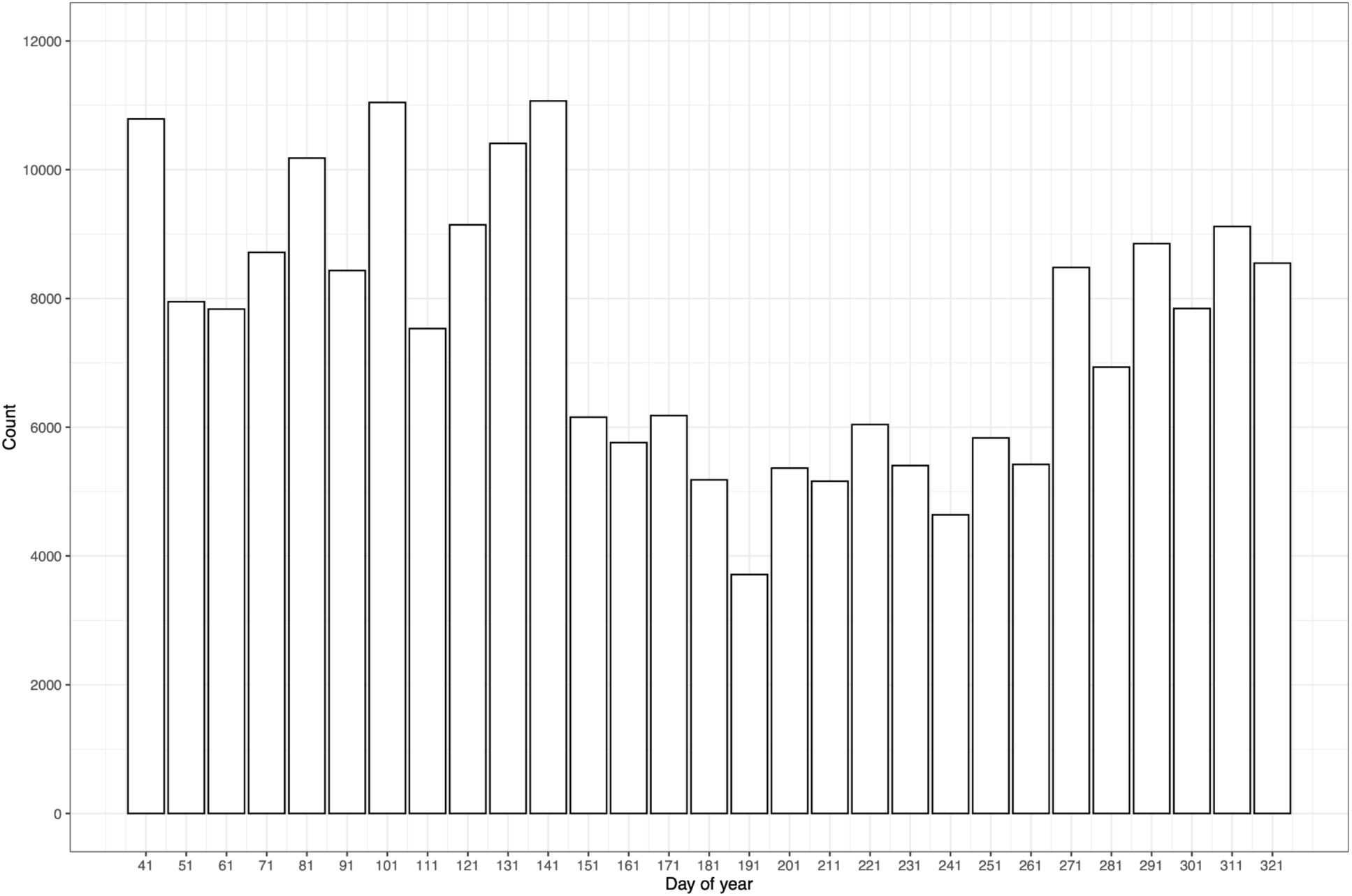
Frequency of missing values that were forward filled. Count is calculated across all samples and years.

#### Model

Using the CONUS field data and the temporal sequence of satellite data described above, we employed a temporal one-dimensional convolutional neural network (1D CNN) architecture, generally following the approach outlined in Allred et al.^21^, with modifications to the temporal sequence and the number of units in certain layers. Fractional cover and canopy gap size class, as measured by field data collection, were modeled together as multi-outputs. As this is an empirical model, we are not directly resolving fractional cover and gap characteristics; rather, we are training the model to learn the empirical relationships among spectral, phenological, and fractional cover/canopy gap characteristics. The input features consisted of the log normalized reflectance values, reflectance vegetation indices, and spatial location (X,Y coordinates). We processed reflectance values and vegetation indices through two separate convolutional streams. Each stream had three consecutive 1D convolutional layers with 64 units, a kernel width of three, rectified linear unit (ReLU) activation, and dilation rates increasing exponentially from one to four with each successive layer. A dropout layer with a dropout rate of 0.2 was applied after each convolutional layer. Final convolutional layers were reduced using average pooling and concatenated, along with a dense layer output of spatial location information. This combined vector was passed through a dense layer with 512 units, followed by a dropout layer of rate 0.2. The final layer contained 16 units with linear activation to predict fractional cover estimates and canopy gap size class together as multi-outputs. We trained the model with a loss of mean squared error, a learning rate of 0.0001, an Adam optimizer, and a batch size of 64. Model performance was evaluated with root mean square error (RMSE), mean absolute error (MAE), and the coefficient of determination (r^2^) of the testing set. There were no architectural designs or post-processing steps to ensure that predictions sum to 100. During inference, predictions were clipped from 0 to 100.

### Data Records

Fractional cover and canopy gap estimates were produced for the 17 western states of the United States (Figure 5)^43^. Estimates for common plant functional types and canopy gap (percentages; refer to Table 2) were produced for all western states; estimates for select genera (percentages; refer to Table 2) were produced for UTM zones 10 through 13. Data are available as Cloud Optimized GeoTIFFs at http://rangeland.ntsg.umt.edu/data/rangeland-s2/. Data are distributed as 75×75 km GeoTIFF tiles within each UTM zone. Tiles overlap by 250 m on all sides. For tiles that intersect two UTM zones, pixels outside the reference UTM zone are masked. GeoTIFFs are in the WGS84 coordinate reference system of their relative UTM zone (EPSG:326XX). All GeoTIFFs are stored as 8-bit integers and have a no data value of 255. Dataset size is approximately 2 TB.

**Figure 5.**
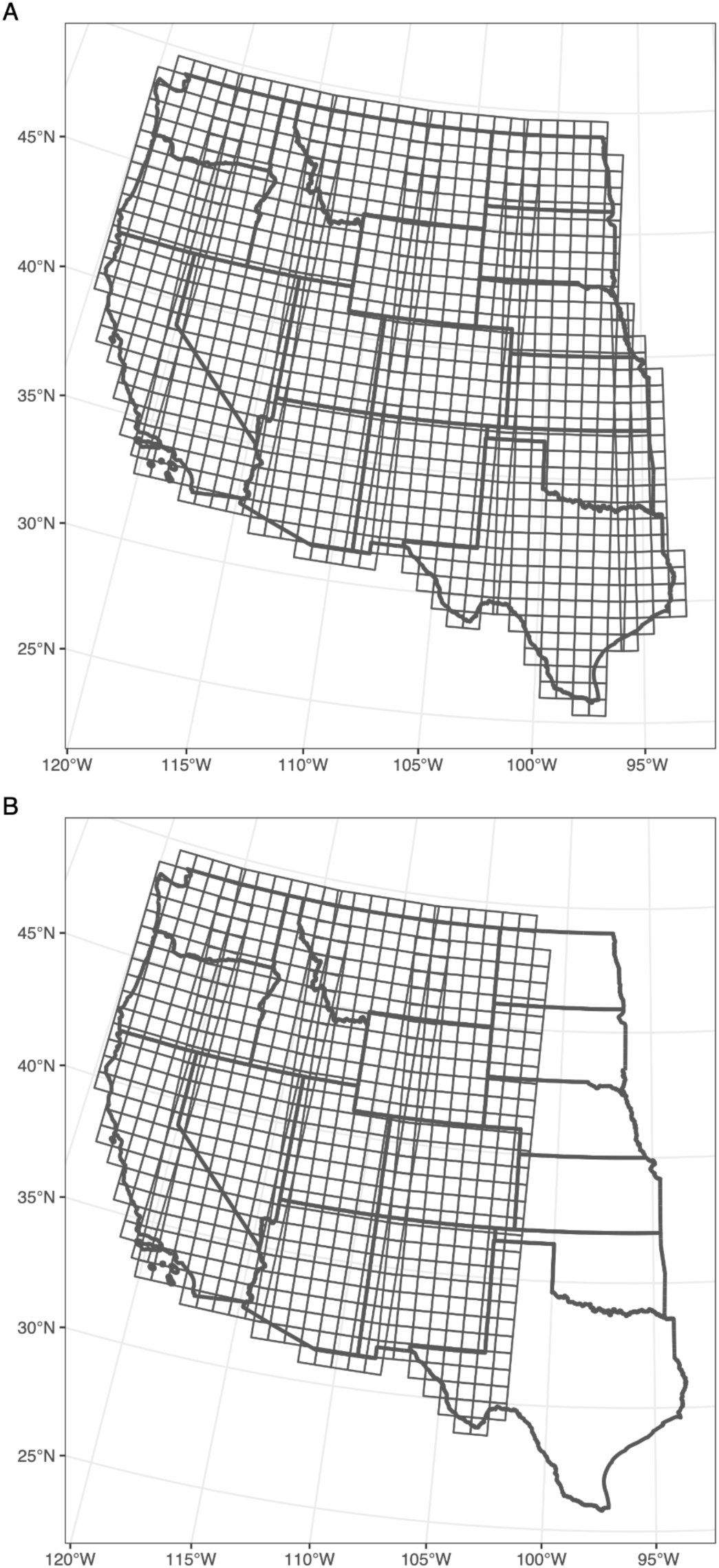
Spatial coverage for A) fractional cover estimates of common plant functional types and canopy gap size class, produced for all western states; and B) fractional cover estimates of select genera sagebrush, invasive annual grass, and pinyon-juniper species, produced for UTM zones 10 through 13. Data were produced for years 2018 through 2024, and are distributed as 75×75 km GeoTIFF tiles within each UTM zone. Refer to Table 2 for definitions of plant categories.

GeoTIFFs are named GRP-YYYY-ZZ-XXXXXX-YYYYYYY.tif, where: GRP = vegetation group; YYYY = four digit year; ZZ = utm zone; XXXXXX = lower left x coordinate; YYYYYYY = lower left y coordinate. Vegetation groups are: 1) pft; fractional cover estimates of common plant functional types; 2) gap; canopy gap estimates of canopy gap size class; 3) arte; fractional cover estimates of sagebrush; 4) iag; fractional cover estimates of invasive annual grass; 5) pj; fractional cover estimates of pinyon-juniper species. GeoTIFFs are organized into top level folders for each vegetation group.

### Technical Validation

#### Model performance

We produced a robust model for rangeland fractional cover and canopy gap sizes. Performance metrics revealed good model fit and strong relationships between predicted and observed values (Table 3; Figures 6 and 7). Overall model performance was similar to or better than Landsat-based predictions of the same plant functional types from the Rangeland Analysis Platform cover dataset (v3)^44^. RMSE of annual forb and grass, perennial forb and grass, shrub, and tree decreased 1.6, 1.3, 1.0, and 0.4 percent, respectively; MAE of the same groups decreased 1.6, 2.2, 1.2, and 0.3 percent, respectively. Relative to an independent evaluation of the same dataset^45^, RMSE of annual forb and grass, perennial forb and grass, shrub, and tree decreased 1.3, 1.8, 1.9, and 1.0 percent, respectively; MAE of the same groups decreased 1.3, 2.0, 2.0, and 0.7 percent, respectively. It is important to note that the reported model error metrics for the Rangeland Analysis Platform cover dataset were produced using a separate testing dataset that included additional years prior to 2018. RMSE of canopy gap size classes 25-50 cm and >200 cm decreased by 0.9 and 2.2, respectively, compared to available Landsat-based predictions^18^.

**Figure 6.**
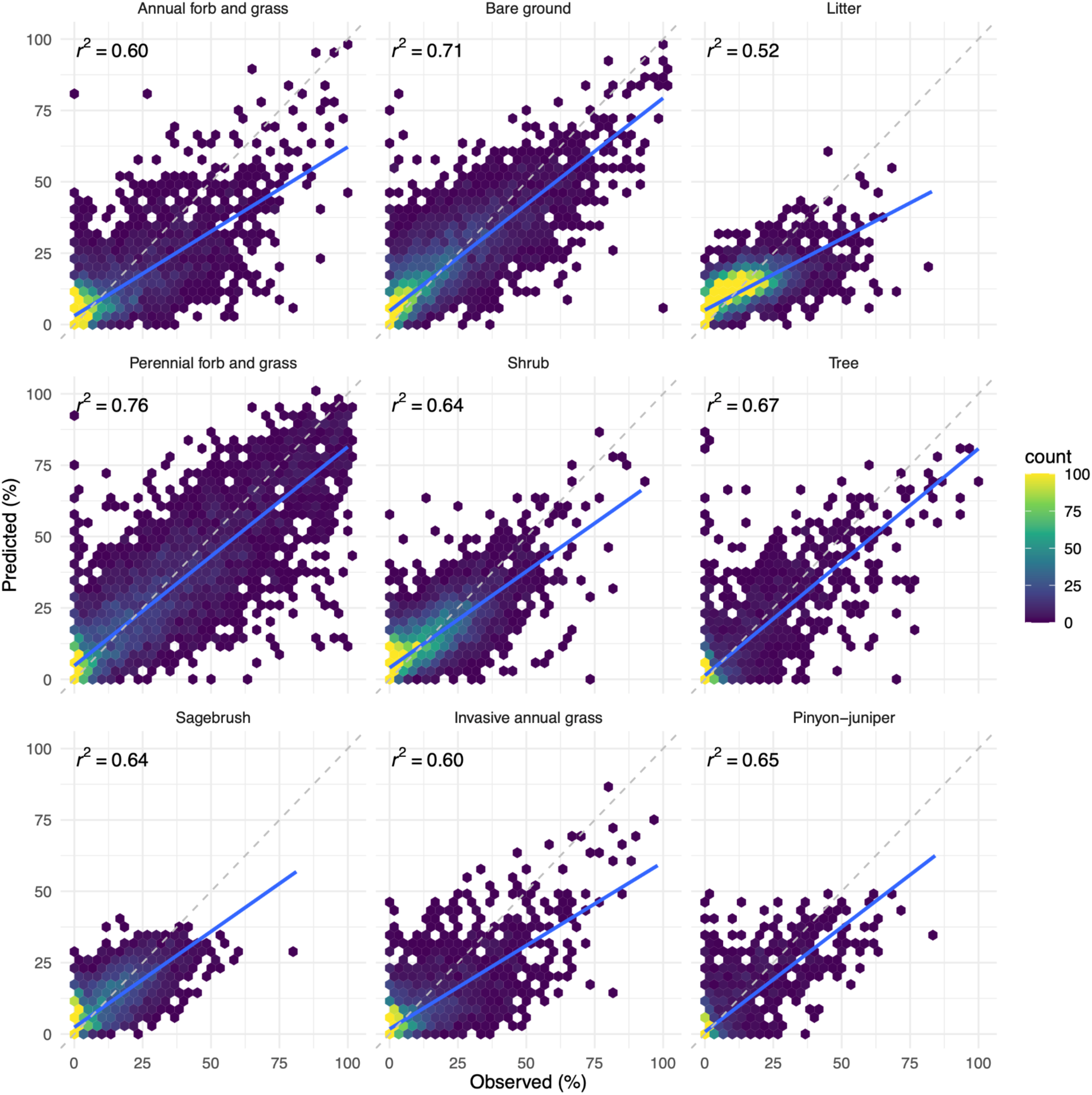
Predictions of fractional cover relative to observed for the testing set. The dashed gray line represents a 1:1 relationship; the solid blue line is the linear fit between predicted and observed. Refer to Table 3 for RMSE and MAE metrics.

**Figure 7.**
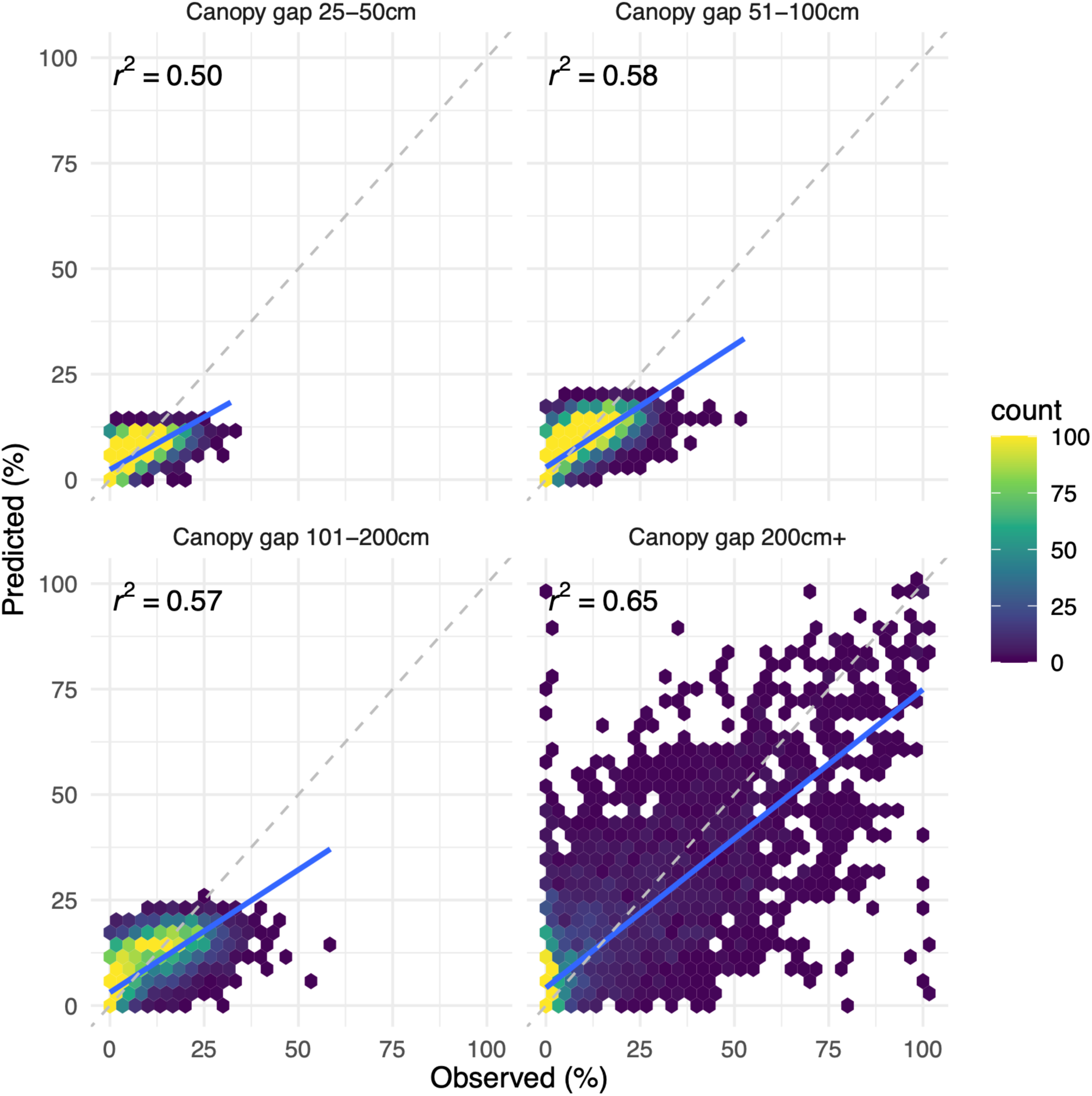
Predictions of canopy gap relative to observed for the testing set. The dashed gray line represents a 1:1 relationship; the solid blue line is the linear fit between predicted and observed. Refer to Table 3 for RMSE and MAE metrics.

**Table 3.**
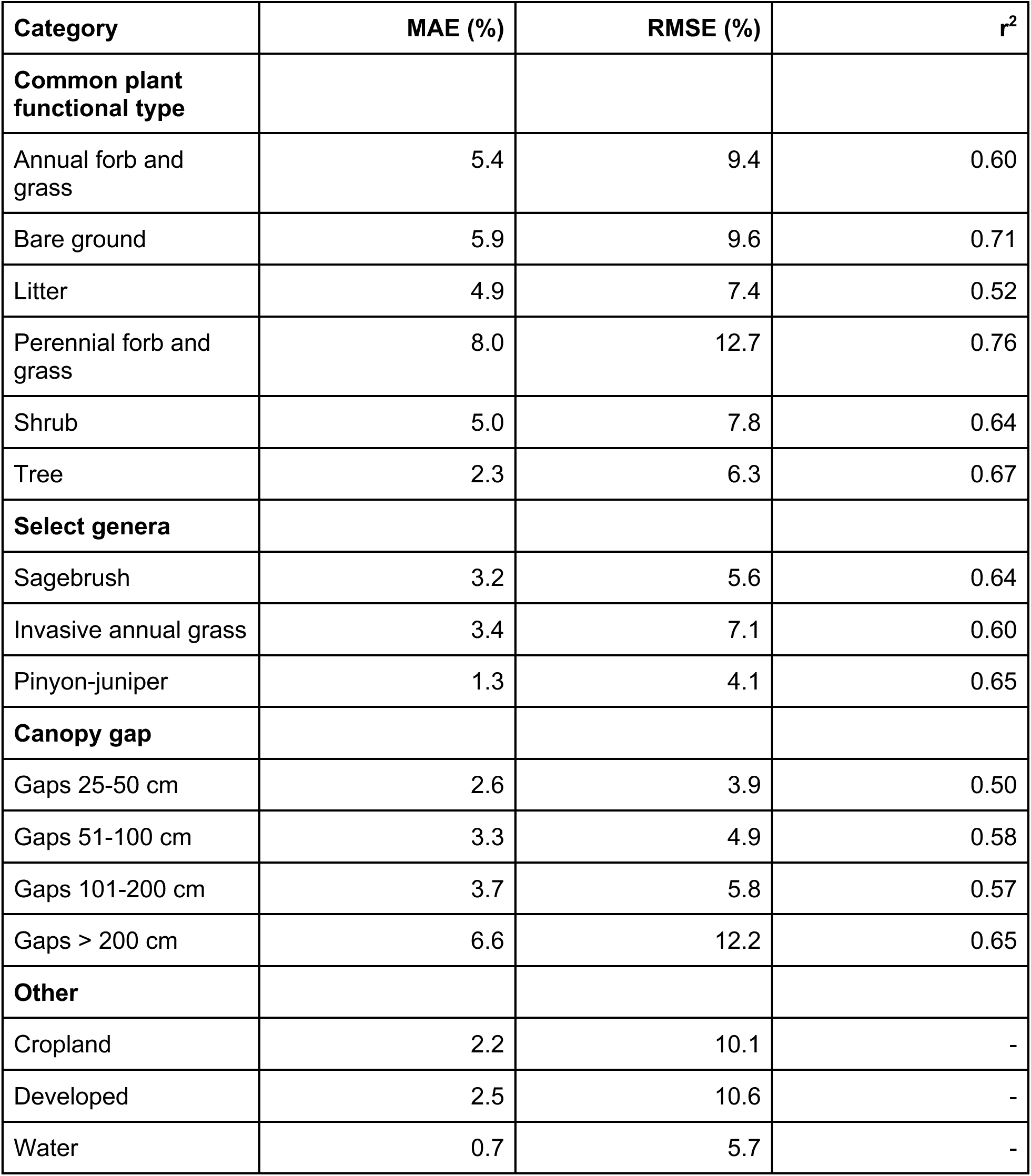
Model performance metrics (mean absolute error, MAE; root mean squared error, RMSE; coefficient of determination, r^2^). Metrics were calculated using testing data (10%) held out from training and validation. Coefficients of determination were omitted for cropland, developed, and water as there were only two possible observed values and provided little relevance.

### Usage Notes

A key strength of Sentinel-2 lies in its ability to capture finer scale spatiotemporal heterogeneity. The 10 m spatial resolution allows for improved detection of small yet ecologically significant features, such as subtle changes in vegetation composition, isolated shrub or tree patches, or management actions. These details, which are often lost at coarser resolutions, can help inform condition assessments and management decisions. For example, managers can use improved and finer resolution estimates to evaluate the effectiveness of invasive species treatments, reclamation efforts, and other management actions^46,47^. Moreover, the more frequent revisit time of Sentinel-2 satellites provides greater temporal resolution, enhancing the ability of the model to use phenology and other temporal dynamics in distinguishing vegetation characteristics. This may be particularly important for vegetation groups such as invasive annual grasses, encroaching coniferous trees, or shrubs which often have very different phenologies compared to surrounding vegetation. As a result, the fractional cover dataset produced (refer to *Data Records*) has more specific plant functional types that can better characterize rangeland vegetation (Figures 8 and 9).

**Figure 8.**
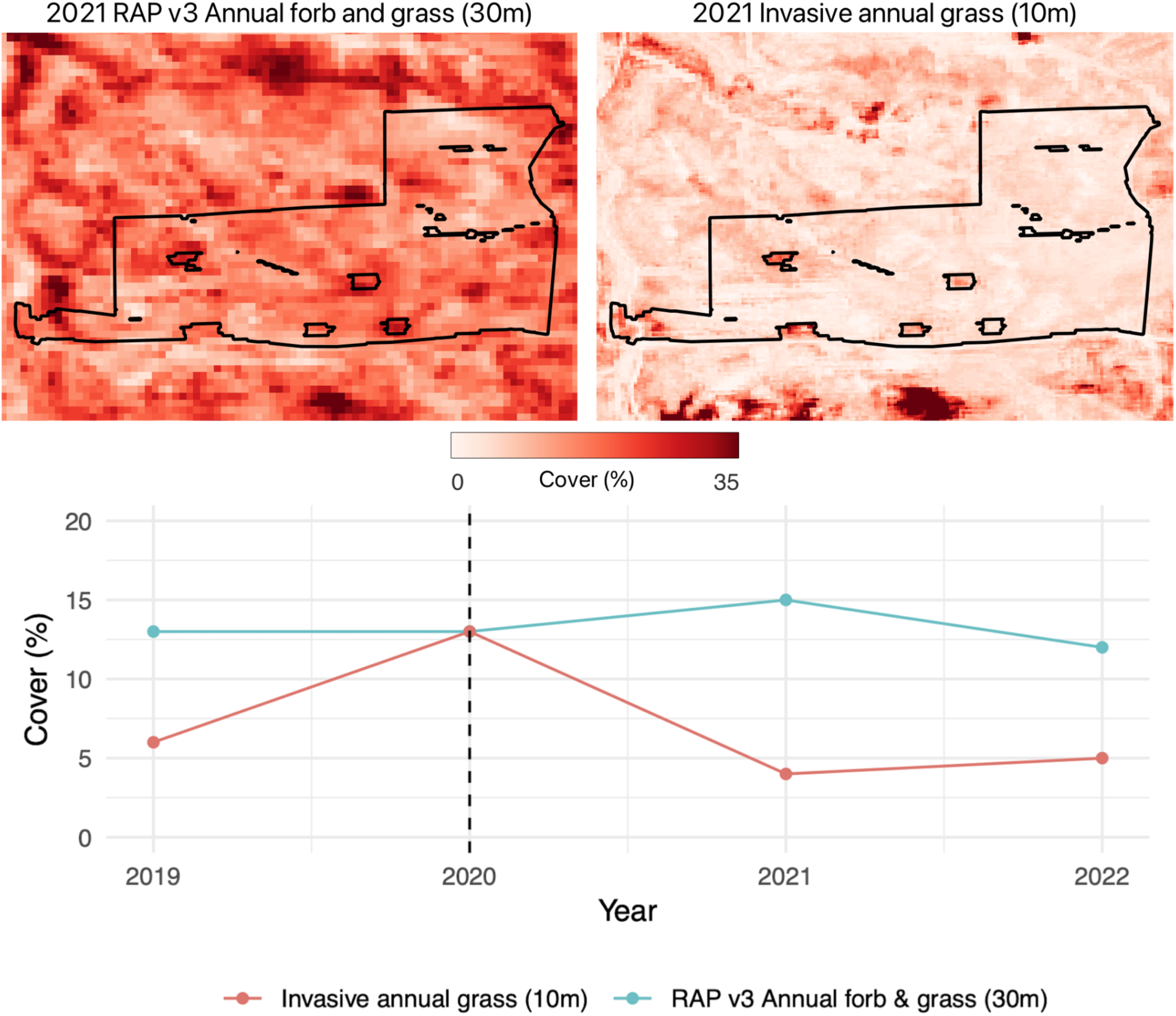
Comparing remotely sensed estimates of Landsat-based fractional cover for annual forb and grass (RAP v3, 30 m resolution)^21^ and Sentinel-2 based invasive annual grass (10 m resolution, described in this paper) for herbicide treatment of invasive annual grasses. Top panels display estimated fractional cover one year post herbicide treatment. Time series represents the average across the treatment for each year; dashed line indicates the year of treatment. The 10 m, Sentinel-2 based estimate of fractional cover detects immediate reductions of invasive annual grasses and correctly highlights areas that were not treated (small inholdings within the treatment). Although cover values do not reach zero, the Sentinel-2 based estimate more closely reflects the near total eradication of invasive annual grasses post treatment (Mealor, written communication, February 21, 2025). Treatment is near Sheridan, Wyoming, USA and is approximately 97 ha.

**Figure 9.**
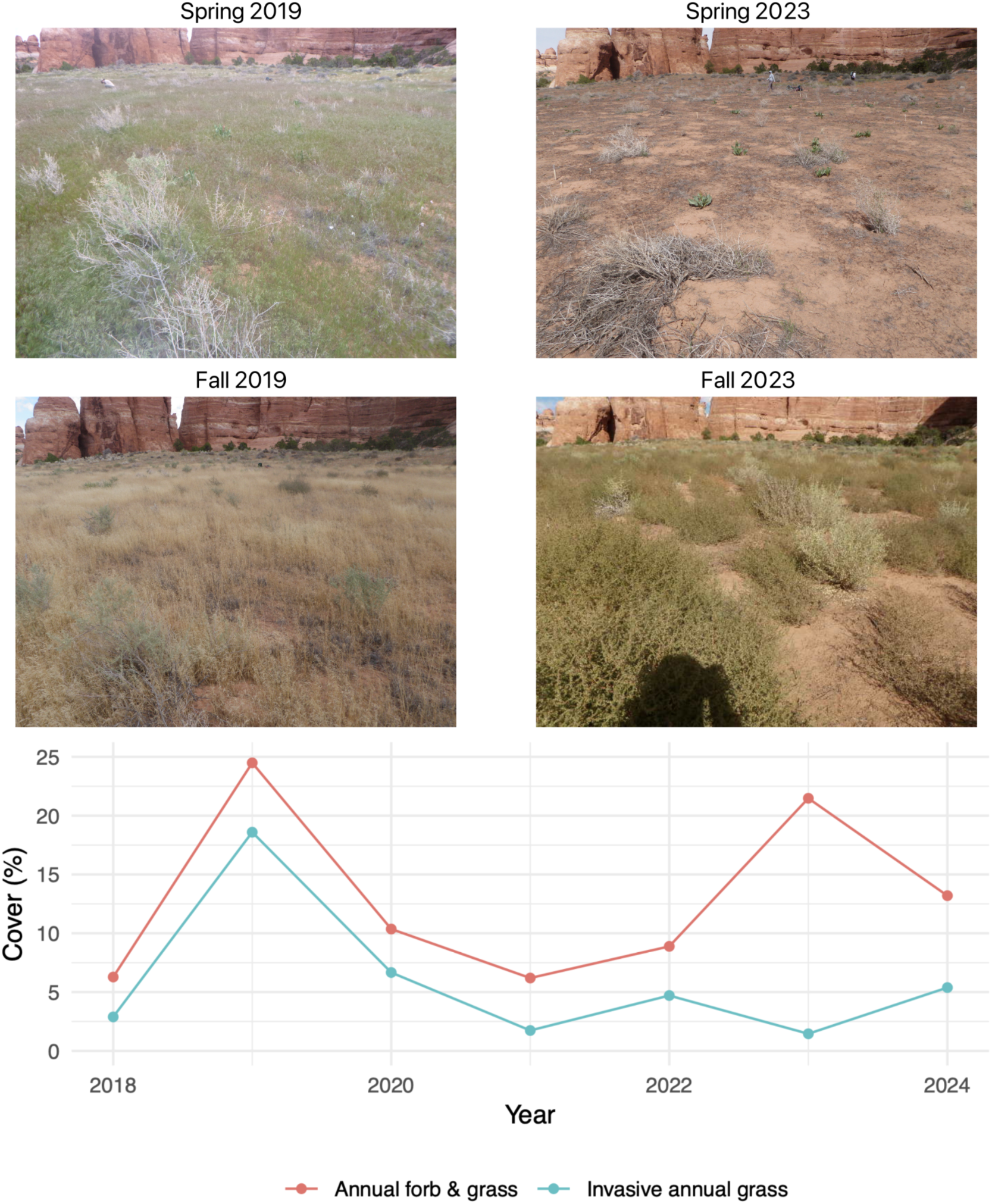
The increased temporal resolution of Sentinel-2 satellites allows the fractional cover model to distinguish between similar plant types with different phenological patterns. The four photographs were taken from the same location in Canyonlands National Park, UT, USA. Spring and fall 2019 photographs show high cover of *Bromus tectorum*, an invasive annual grass species that completes its life cycle during the spring and early summer. Spring and fall 2023 photographs show the same location in a year dominated by *Salsola tragus*, an introduced annual forb that reaches peak biomass in mid- to late summer. The predicted cover for this location shows high cover of annual forb and grass in both of the years pictured, but correctly estimates high cover of invasive annual grass species only in 2019. Photo credit: U.S. Geological Survey.

#### Important considerations

While this model represents an advancement in estimating fractional cover and canopy gap, users should understand limitations in order to employ the data responsibly. These considerations are not shortcomings of the model itself but rather factors to account for when interpreting and applying the data.

The Sentinel-2 imagery used in this model covers a relatively recent time period. For analyses requiring long-term historical trends, datasets based on Landsat imagery^18,21,48^ may be more appropriate. While harmonization between Sentinel-2 and Landsat-derived cover estimates (i.e., harmonization between the data described here and cover estimates derived from Landsat) can be an additional area of research, the exploration and testing of such work has yet to be completed.

While the model demonstrates strong overall performance, certain vegetation groups presented unique challenges. Understanding these challenges is vital for accurate interpretation.

Although we produced these data across a broad region of the western United States, they are primarily intended for rangeland ecosystems. Cover estimates may not perform well in ecosystems that do not share the same vegetation or canopy gap characteristics as rangelands.

Distinguishing sagebrush from other shrub species, particularly rabbitbrush (*Ericameria nauseosa*) in the Great Basin, proved challenging, sometimes leading to overestimation of sagebrush (refer to Figure 5). Conversely, we noticed good discrimination between sagebrush and greasewood (*Sarcobatus vermiculatus*) in the Great Basin, whereas in the Northern Great Plains, the model struggled to differentiate sagebrush from greasewood, particularly in riparian areas. Critically, sagebrush estimates are only reliable within the known sagebrush biome^49,50^; estimates outside this range should be interpreted with caution and considered uncertain.

Invasive annual grasses often have unique characteristics compared to native annual or perennial grasses that lend themselves to identification with satellite remote sensing, for example earlier green up and senescence, multiple germinations or die-off, and increased response to precipitation^51–53^. Care should be taken, however, to corroborate that native annual forbs and grasses are not being misidentified as invasive. For example, in very dry years in the Great Plains, the native annual grass *Vulpia octoflora* can exhibit similar growth patterns as invasive annual grasses, and we noticed that large patches of *V. octoflora* were sometimes categorized as invasive annual grass. Comparing predictions of invasive annual grass cover to annual forb and grass cover, and to local expertise and knowledge, may help mitigate misidentification.

Pinyon-juniper woodlands are notoriously difficult to characterize with satellite data^54–56^. The model captures general spatiotemporal trends in pinyon-juniper cover, but the estimates should not be interpreted as a definitive presence/absence of pinyon-juniper. Therefore, estimates are only valid within established pinyon-juniper ranges^57^. Predictions of pinyon-juniper cover in dry forests, semidesert shrublands, or ecotones between these ecosystems and pinyon-juniper woodlands should be treated with caution; for example, the model estimates cover of pinyon-juniper throughout the dry forests of the central Sierra Nevada where pinyon-juniper is generally absent.

Due to the model’s methodology and species aggregation (refer to Table 1), understory herbaceous or woody vegetation may not be well-represented, especially in areas with dense overstory canopies (e.g., southern Texas Tamaulipan thornscrub, Rocky Mountain forests).

Additionally, users should be aware that woody vegetation groups (shrubs and trees) may exhibit year-to-year variability, between or within groups, potentially due to noise. Thus it may be helpful for the user to examine and combine woody vegetation groups to better understand variability.

Riparian and wetland areas also presented unique challenges. Mischaracterization of vegetation (e.g., attributing herbaceous cover to trees or shrubs) and nonsensical estimates (e.g., attributing vegetated areas to developed) were observed in some cases. These challenges are partly due to riparian and wetland areas exhibiting unique phenological patterns compared to upland ecosystems, with different timing of green-up, senescence, and vegetative responses to seasonal changes^58^. Because these ecosystems are often under-sampled in field surveys, the model’s characterization of their dynamics may be limited. Careful inspection of the data in these areas, informed by local expertise, is essential.

#### Best practices

These considerations highlight the importance of using these data in conjunction with other sources, including local knowledge, field data, and other relevant information. Integrating diverse data sources is crucial for accurate interpretation and informed decision-making^59^. By understanding the model’s capabilities and limitations, and by applying local expertise, users can effectively leverage these data for rangeland management and research.

## Code and Data Availability

The model architecture and weights are available on GitHub (https://github.com/allredbw/rangeland-s2-cover-gap).

Data are available as Cloud Optimized GeoTIFFs at http://rangeland.ntsg.umt.edu/data/rangeland-s2/

## Acknowledgements

We thank the many field data collectors and providers who shared their data with us. We thank Megan Creutzburg, Carrie-Ann Houdeshell, and Jeremy Maestas for testing and intellectual guidance. This research used computer resources provided by 1) the SCINet project and/or the AI Center of Excellence of the USDA Agricultural Research Service, ARS project numbers 0201-88888-003-000D and 0201-88888-002-000D; 2) Google Earth Engine; and 3) the Numerical Terradynamic Simulation Group at the University of Montana. Views expressed in this article are those of the authors and do not necessarily represent views of the U.S. Fish and Wildlife Service or any official USDA determination or policy, but do represent the views of the U.S. Geological Survey. Any use of trade, firm, or product names is for descriptive purposes only and does not imply endorsement by the U.S. Government.

## Author information

### Contributions

Allred performed data retrieval, modeling, testing, inference, and wrote the manuscript. McCord performed data retrieval, testing, and wrote the manuscript. Morford performed testing and wrote the manuscript. Assal, Bestelmeyer, Boyd, Brooks, Cady, Duniway, Fuhlendorf, Green, Harrison, Jensen, Kachergis, Knight, Mattilio, Mealor, Morford, Naugle, O’Leary, Olsoy, Peirce, Reinhardt, Shriver, Smith, Tack, A. Tanner, E. Tanner, Twidwell, and Webb performed testing, assisted with the manuscript, and provided intellectual guidance.

